# Multiple cyanotoxin congeners produced by sub-dominant cyanobacterial taxa in riverine cyanobacterial and algal mats

**DOI:** 10.1101/705848

**Authors:** Laura T. Kelly, Keith Bouma-Gregson, Jonathan Puddick, Rich Fadness, Ken G. Ryan, Timothy W. Davis, Susanna A. Wood

## Abstract

Benthic cyanobacterial proliferations in rivers are have been reported with increasing frequency worldwide. In the Eel and Russian rivers of California, more than a dozen dog deaths have been attributed to cyanotoxin toxicosis since 2000. Periphyton proliferations in these rivers comprise multiple cyanobacterial taxa capable of cyanotoxin production, hence there is uncertainty regarding which taxa are producing toxins. In this study, periphyton samples dominated by the cyanobacterial genera *Anabaena* spp. and *Microcoleus* spp. and the green alga *Cladophora glomerata* were collected from four sites in the Eel River catchment and one site in the Russian River. Samples were analysed for potential cyanotoxin producers using polymerase chain reaction (PCR) in concert with Sanger sequencing. Cyanotoxin concentrations were measured using liquid chromatography tandem-mass spectrometry, and anatoxin quota determined using droplet digital PCR. Sequencing indicated *Microcoleus* sp. and *Nodularia* sp. were the putative producers of anatoxins and nodularins, respectively, regardless of the dominant taxa in the mat. Anatoxin concentrations in the mat samples varied from 0.1 to 18.6 μg g^−1^ and were significantly different among sites (*p* < 0.01, Wilcoxon test); however, anatoxin quotas were less variable (< 5-fold). Dihydroanatoxin-a was generally the most abundant variant in samples comprising 38% to 71% of the total anatoxins measured. Mats dominated by the green alga *C. glomerata* contained both anatoxins and nodularin-R at concentrations similar to those of cyanobacteria-dominated mats. This highlights that even when cyanobacteria are not the dominant taxa in periphyton, these mats may still pose a serious health risk and indicates that more widespread monitoring of all mats in a river are necessary.

## 1. Introduction

Reports of toxic benthic cyanobacterial proliferations have been described over the past 30 years [e.g. 1, 2, 3] and are now increasing in frequency globally [4–9]. Despite these reports, investigations into benthic cyanotoxin producers, and variability in toxin production are limited [7]. Many benthic cyanobacteria are now known to produce cyanotoxins, for example, *Anabaena, Phormidium*, *Lyngbya*, *Oscillatoria*, *Nostoc*, *Nodularia* and *Microcoleus*. While these taxa are often dominant in proliferations [7], they are also common components of periphyton mats where other algae are more abundant [e.g. *Cladophora glomerata*; 10, 11], and the risks these mats pose is relatively unknown.

Cyanotoxins are typically classified by their different toxicological properties into neurotoxins (e.g., anatoxins and saxitoxins [STXs; 12], hepatotoxins (e.g., microcystins [MCYs], nodularins [NODs] and cylindrospermopsins [CYNs]), cytotoxins and dermatotoxins [13, 14]. Among benthic cyanobacteria, anatoxins are the most commonly reported cyanotoxin. Anatoxins comprise four main structural congeners; anatoxin-a (ATX), dihydroanatoxin-a (dhATX), homoanatoxin-a (HTX) and dihydrohomoanatoxin-a (dhHTX), and their relative proportions vary in environmental samples [15].

Recently toxic benthic cyanobacteria have been recorded in Californian streams (Fetscher et al. 2015; Bouma-Gregson et al. 2018; Anderson et al. 2018). Anatoxins, MCYs, NODs and CYNs have all been recorded in the mats and in samples collected using solid-phase absorption toxin tracking (SPATT) samplers [16, 17]. A number of sites on the Eel and Russian rivers in Northern California regularly experience benthic cyanobacterial and chlorophyte proliferations [16]. The mats at these sites contain assemblages comprised of a variety of cyanobacterial genera known to produce cyanotoxins. Extensive mats dominated by the green alga *C. glomerata*, are often also present at these locations, with low levels of cyanobacteria in the mats. There is uncertainty about which genera in these systems are producing toxins, although *Microcoleus* has been identified as an anatoxin producer in the Eel River using assembled metagenomes [18], and cultures of *Microcoleus* isolated from the Russian River also produce anatoxins [19]. Both toxic and non-toxic genotypes occur in the genus *Microcoleus*, and these genotypes can be present at varying relative abundances in environmental mats [15, 20], resulting in considerable spatial variability in anatoxin concentrations of mats within short distances.

Identifying cyanotoxin producers with culture-based studies can be prohibitively costly considering the range of potential toxic taxa present and the spatial variability in toxin production; however, molecular techniques (e.g., PCR and quantitative PCR) can be used to screen for the presence of genes involved in cyanotoxin biosynthesis. These techniques, coupled with DNA sequencing, can provide a strong indication of the toxin producers in environmental samples and this information can then be used to guide culturing of selected species. Molecular assays to determine the concentration of *anaC* gene copies using quantitative PCR (qPCR) and droplet digital PCR (ddPCR) have recently been developed and allow the toxin quota (amount of toxin per toxic cell) to be determined [15, 20]. Comparisons of toxin quota, combined with data on the physicochemical conditions at the time of sampling, may provide insights into the drivers of anatoxin production.

To gain a better understanding of the diversity of cyanotoxins and cyanotoxin producers in the Eel and Russian rivers of Northern California, this study aimed to: (1) identify potential benthic cyanotoxin producers, (2) assess spatial variability in anatoxin concentrations and quotas within and between mats, sites and rivers, and (3) determine cyanotoxin levels in *Cladophora*-dominated mats. Periphyton samples were collected from the Eel and Russian rivers. PCR amplification of genes involved in toxin production combined with DNA sequencing of the amplified genes were used to identify potential cyanotoxin producers and liquid chromatography tandem-mass spectroscopy (LC-MS/MS) was used to quantify anatoxin congeners, nodularin-R (NOD-R) and MCYs. Spatial variability in anatoxin concentrations and anatoxin quotas were investigated by combining toxin concentrations determined by LC-MS/MS with ddPCR quantification of *anaC* gene copies.

## 2. Materials and Methods

### 2.1 Environmental sample collection

Sampling sites were chosen in the Eel and Russian rivers in Northern California. The riverbeds at the sampling sites were comprised of gravel, cobble or boulders. The four sites in the Eel River watershed were sampled on 29 July 2018 and spanned upstream drainage areas of 17 km^2^ to 495 km^2^. Site 1_ELD was on Elder Creek and sites 2_SFE – 4_SFE were on the South Fork Eel River (Fig 1). The single Russian River site (5_RUS) was sampled on 31 July 2018 and had an upstream drainage area of 793 km^2^ (Fig 1). At each site, ten samples with visible cyanobacterial mats were selected and attached cyanobacteria collected by scraping a small sample (ca. 2 cm diameter) into a 15 mL centrifuge tube. At 4_SFE, fine-scale samples were collected by taking five samples from each of three additional rocks. Additional samples of *C. glomerata* mat material were collected at 3_SFE on the South Fork Eel River. Samples were stored on ice, frozen (–20 °C) on return to the laboratory (within 8 hours), and subsequently lyophilised prior to further analysis. At each site dissolved oxygen, temperature, pH, and conductivity were measured with handheld probes. Filtered water samples were also collected (0.7 μm Whatman GF/F) and dissolved inorganic nitrogen and dissolved reactive phosphorus were measured using EPA methods 300.1, 351.2 and 365.2.

**Fig 1.**
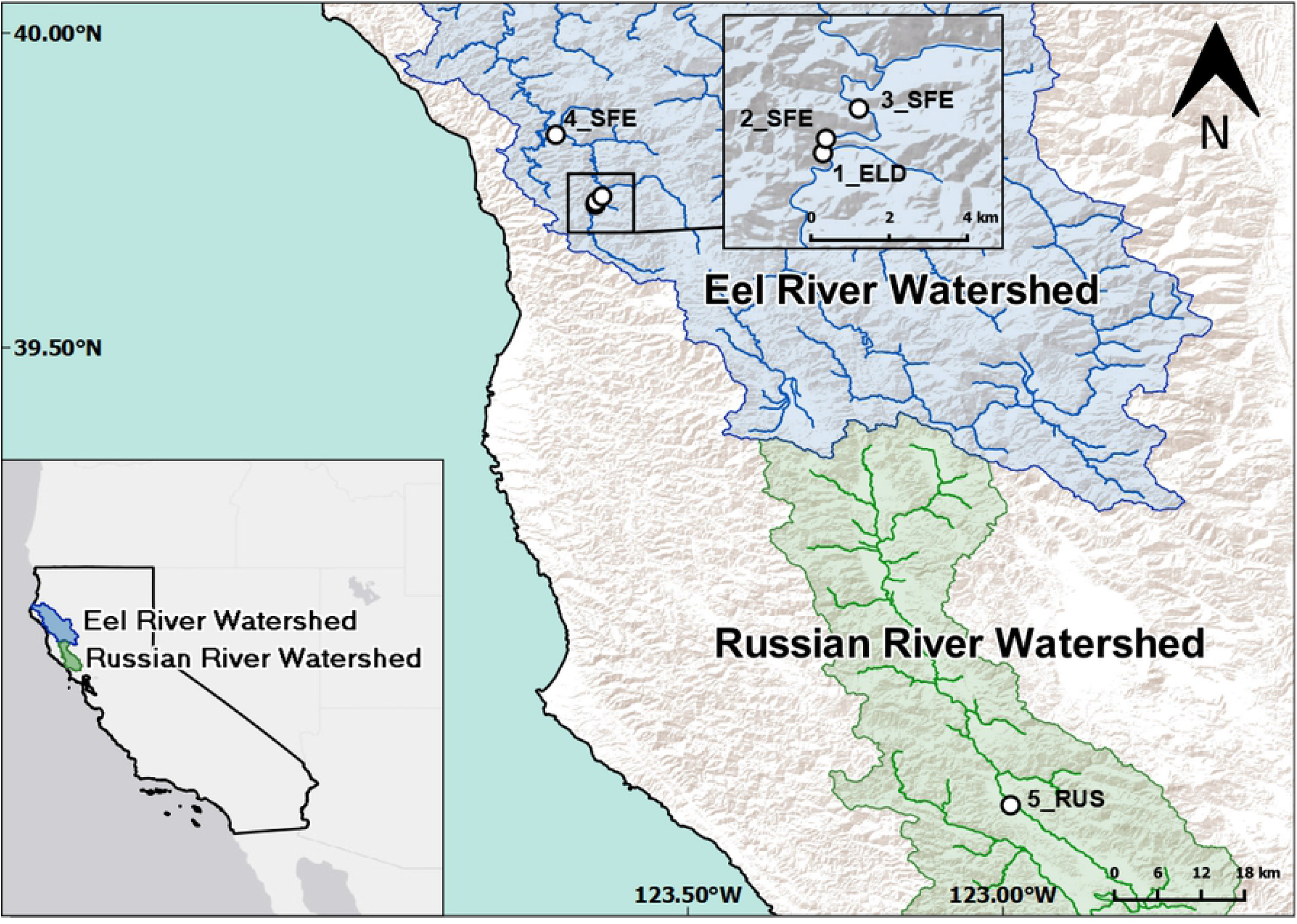
Map of sampling sites on the South Fork Eel River and on the Russian River in California, USA. Blue areas on the map refer to the Eel River watershed, while green denotes the Russian River watershed. Sites are labelled in order of increasing drainage area from 1 to 5, while the suffix refers to the river: ELD = Elder Creek, SFE = South Fork Eel, RUS = Russian River.

All ten samples from each site were analysed for cyanotoxin concentrations. Three samples from each site were screened for cyanotoxin genes, which were sequenced if detected. Anatoxin quotas (the amount of anatoxin per toxigenic cell) were determined for the five samples with the highest anatoxin concentrations from sites 3_SFE, 4_SFE and 5_RUS, as well as the fine-scale samples from 4_SFE (S1 Fig). These sites were selected because anatoxin concentrations were markedly higher at these locations, and practical constraints limited the number of samples that could be processed.

### 2.2 Toxin analysis

Lyophilised environmental samples were homogenised with a sterile metal spatula and sub-sampled for cyanotoxin analysis by suspending ca. 10 mg dry weight of mat material in 1 mL of 0.1% formic acid in water. The samples were frozen at –20 °C and subsequently thawed in a bath sonicator for 30 min (53 kHz; LHC bath sonicator, Kudos, Shanghai, China). Freeze-thaw and sonication steps were repeated two more times for each sample. Cyanotoxin extracts were clarified by centrifugation (14,000 × *g*, 5 mins) and the supernatants transferred to septum-capped glass vials for analysis of anatoxins, CYNs, MCYs and NOD-R using liquid chromatography tandem-mass spectrometry (LC-MS/MS).

Anatoxins (ATX, HTX, dhATX and dhHTX) and CYNs (CYN and deoxycylindrospermopsin; doCYN) were analysed using the LC-MS/MS methodology described in Wood et al. [21]. Multiple-reaction monitoring (MRM) channel values for the anatoxins are provided in Wood et al. [21] and quantitation MRM channels for CYN and doCYN were 416.3 > 194.15 and 400.3 > 194.15, respectively. The anatoxins and cylindrospermopsins were quantified using a mixed external five-point calibration curve (0.5 ng mL^−1^ – 18 ng mL^−1^ in 0.1% formic acid) made from certified reference materials for ATX and CYN (National Research Council, Canada) and a standard for dhATX calibrated by quantitative nuclear magnetic resonance spectroscopy. The concentrations of ATX and HTX were determined using the ATX calibration curve and the concentrations of dhATX and dhHTX were determined using the dhATX calibration curve and the concentration of CYN and doCYN were determined using the CYN standard. The analytical limit of detection (LoD) for ATX, dhATX and CYN was 0.05 ng mL^−1^, which equates to an approximate LoD of 0.005 μg g^−1^ in the sample extracts (dependent on the amount of sample weighed out for extraction). The limit of quantitation (LoQ) was 0.02 μg g^−1^ dw.

Microcystins and NOD-R were analysed as described in Wood et al. [22] using a mixed external four-point calibration curve (2 ng mL^−1^ – 100 ng mL^−1^ in 50% methanol) made up of standards for NOD-R, MCY-RR, -YR and -LR (DHI Lab Products, Denmark). The analytical LoD for MCY-RR, -YR, -LR and NOD-R was 0.02 ng mL^−1^, which equates to an approximate LoD of 0.002 μg g^−1^ in the sample extracts (dependent on the amount of sample weighed out for extraction). The LoQ was 0.006 μg g^−1^ dw.

### 2.3 Molecular analyses

A sub-sample (ca. 15 mg dry weight) was placed into the first tube of a PowerSoil® DNA Isolation Kit (Qiagen, CA, USA) and DNA extracted according to the manufacturer’s protocols. The extracted DNA was quantified (NP80 NanoPhotometer, Implen GmbH, Munich, Germany) and stored at –20 °C until further molecular analyses.

Samples were screened by PCR (S2 Table) for the presence of *mcyE/ndaF* [primers HEPF/HEPR; 23], *sxtA* [primers Sxta/Sxtf; 24], *anaC* [primers anaC-gen-F/anaC-gen-R; 25] and *cyl* [primers cynsufF/cylnamR; 26]. The reactions consisted of 12.5 μL MyTaq RedMix (Bioline, London, UK), 1 μL each of the relevant forward and reverse primer (S2 Table), 3 μL bovine serum albumin (BSA; Sigma, USA), 4.5 μL DNA/RNA free water (Thermo Fisher Scientific) and 3 μL of template DNA. The cycling conditions for all reactions comprised an initial denaturation at 95 °C for 1 min, followed by 30 cycles with denaturation at 95 °C for 15 s, annealing at 54 °C for 15 s extension at 72 °C for 15 s and a final extension at 72 °C for 5 min and hold at 4 °C.

For sequencing, positive PCR reactions were purified using a Nucleospin PCR clean-up kit (Machery-Nagel, Düren, Germany), according to the manufacturer’s directions. Purified PCR product was then quantified (NP80 NanoPhotometer, Implen GmbH, Munich, Germany) and diluted to a concentration of 5 ng μL^−1^ (*mcyE*) or 4 ng μL^−1^ (*anaC*). Amplicons were sequenced bi-directionally with gene-specific primers using the BigDye Terminator v3.1 Cycle Sequencing Kit (Applied Biosystems, USA). Sequences were compared for similarity to reference sequences using Blastn (NCBI). Sequences obtained in this study were deposited in GenBank under accession numbers MK821061 to MK821086.

The absolute number of copies of *anaC* gene were quantified by ddPCR, using the *Phor-AnaC* primers and probes (S2 Table) according to the methods of Wood and Puddick [15]. Anatoxin quota were calculated as the summed anatoxins per mg dw divided by the *anaC* copy number per mg dw as described by Kelly et al. [20].

### 2.6 Statistical analysis

All statistical analyses were conducted in the software R Studio (R Version 3.5.1). Mean toxin concentrations, *anaC* gene copy numbers and anatoxin quota were tested for normality using the Shapiro-Wilks test. All variables failed to meet parametric test assumptions, so non-parametric tests were used. Comparisons of mean toxin concentrations and anatoxin quota were undertaken using Kruskal-Wallis tests and pairwise Wilcoxon rank sum tests with Benjamini-Hochberg adjustment for multiple comparisons.

## 3. Results

### 3.1 Cyanotoxin presence and variability

All five sites surveyed had visible cyanobacterial biomass present. Sites 1_ELD, 2_SFE and 4_SFE were riffle habitats, and cyanobacterial mats were cohesive, attached to benthic cobbles, and were a black/brown colour characteristic of *Microcoleus-dominated* mats in these rivers. Site 3_SFE was a slow-moving pool and was dominated by senescing *C. glomerata* and spires of *Anabaena* spp., with *Nostoc* spp. also present upstream of the survey site. Site 5_RUS was a run habitat on the Russian River. It was also dominated by *C. glomerata* and epiphytized by *Anabaena* spp. Environmental conditions at the sites were similar (S1 Table). Water column nitrogen concentrations ranged from 0.13 – 0.15 mg L^−1^ and phosphorus concentrations ranged from 0.010 – 0.020 mg L^−1^.

Anatoxins were detected by LC-MS/MS at all five sites, though levels were low (< 0.2 μg g^−1^ dw) at sites 1_ELD and 2_SFE (Fig 2A). Total anatoxin concentrations at the other three sites were considerably higher, with samples ranging up to 18.6 μg g^−1^ dw. The mean anatoxin concentration at sites 3_SFE, 4_SFE and 5_RUS were significantly different both from each other and from 1_ELD and 2_SFE (Fig 2A; *p* < 0.01; Wilcoxon test). Hepatotoxins comprised NOD-R, MCY-LR, and dmMCY-LR and were detected at all sites, though the concentrations in 4_SFE and 5_RUS were very low and only detected in one (4_SFE) or two (5_RUS) samples (Fig 2B). Site 3_SFE had the highest hepatotoxin concentrations (mean 1.4 μg g^−1^ dw), with all other site concentrations between the LOD of 0.002 μg g^−1^ and LOQ of 0.006 μg g^−1^ dw. Microcystin/nodularin concentrations were below quantitation limits at most sites. Mean concentrations (± standard error) of anatoxins in the *C. glomerata* mats were 7.4 μg g^−1^ dw (± 0.15 μg g^−1^ dw) and mean nodularin concentrations were 0.33 μg g^−1^ dw (± 0.27 μg g^−1^ dw).

**Fig 2.**
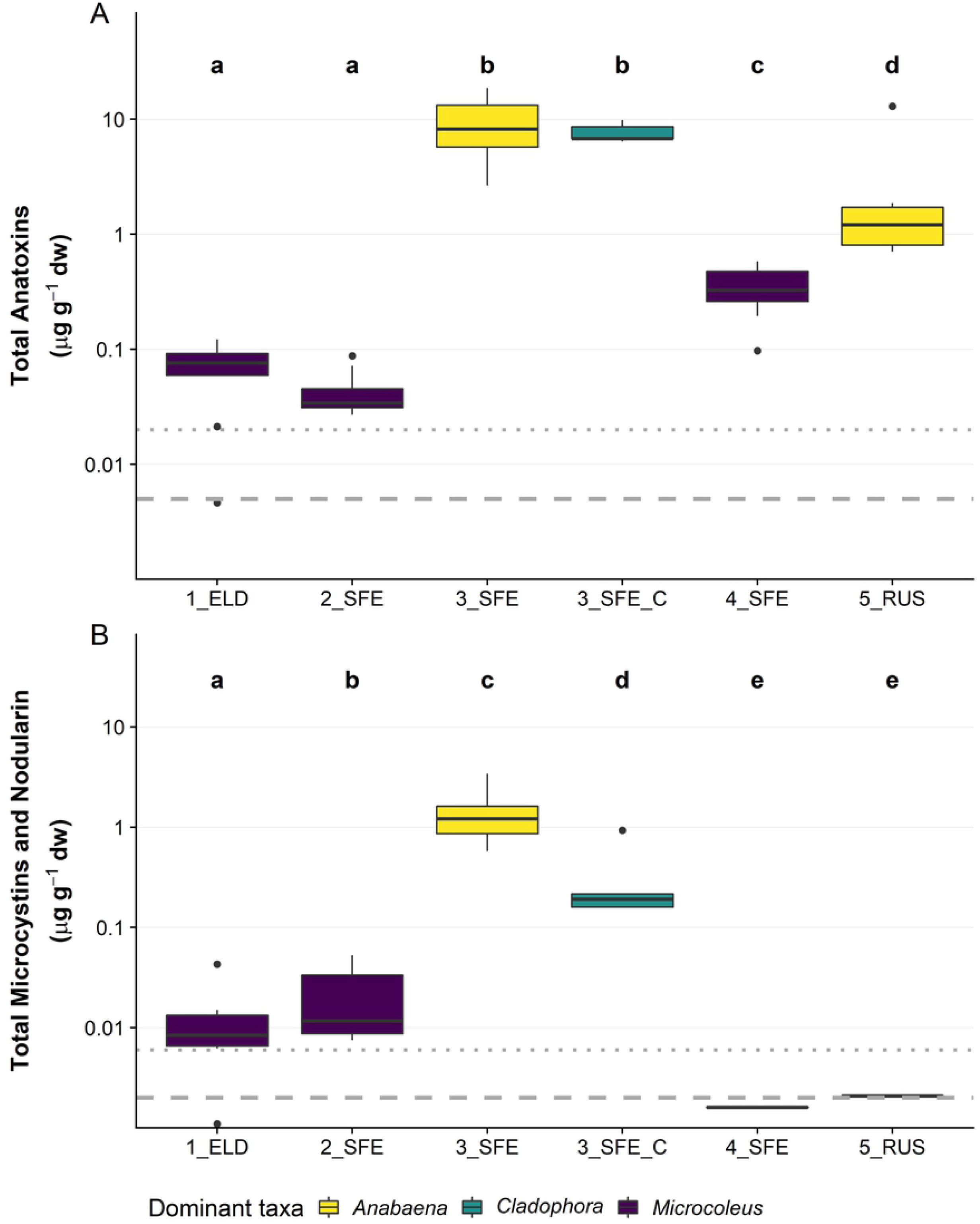
Anatoxin and microcystin/nodularin concentrations vary by site. (A) Summed anatoxin (the total of the four congeners; anatoxin-a, homoanatoxin-a, dihydroanatoxin-a and dihydrohomoanatoxin-a); and (B) summed microcystins and nodularin-R from 10 replicate attached periphyton samples (*n* = 5 for *Cladophora*-dominated samples; 3_SFE_C). collected from each of four sites on the Eel River on 29 July 2018 and 9 attached periphyton samples from one site on the Russian River on 31 July 2018. Colours represent the dominant taxa in mats collected from each site. dw = dry weight. Note log scale on the *y*-axes. Lines within the boxes are medians, the ends of boxes are quartiles and whiskers extend to the lowest or highest data point ≤ 1.5 × the interquartile range. Black dots are outliers. A Kruskal-Wallis test and pairwise Wilcoxon rank sum test with a Benjamini-Hochberg adjustment was used to identify sites that were significantly different from one another (p <0.05), denoted by the letter above the plot. Dotted lines represent the analytical limit of quantitation and bold dashed lines the analytical limit of detection.

The anatoxins detected in the samples were almost exclusively comprised of ATX and dhATX, with low levels (< 0.01 μg g^−1^ dw) of HTX and dhHTX only detected at 1_ELD and 2_SFE (Fig 3). The composition of anatoxin congeners was similar within each site (Table 1), with the exception of 4_SFE, which had more variable proportions of ATX and dhATX among samples. Hepatotoxins (MCYs and NODs) were almost exclusively comprised of NOD-R, with only low levels (< 0.01 μg g^−1^ dw) of MCY-LR and dmMCY-LR detected. Nodularin-R was the only hepatotoxin detected at 1_ELD and 5_RUS.

**Fig 3.**
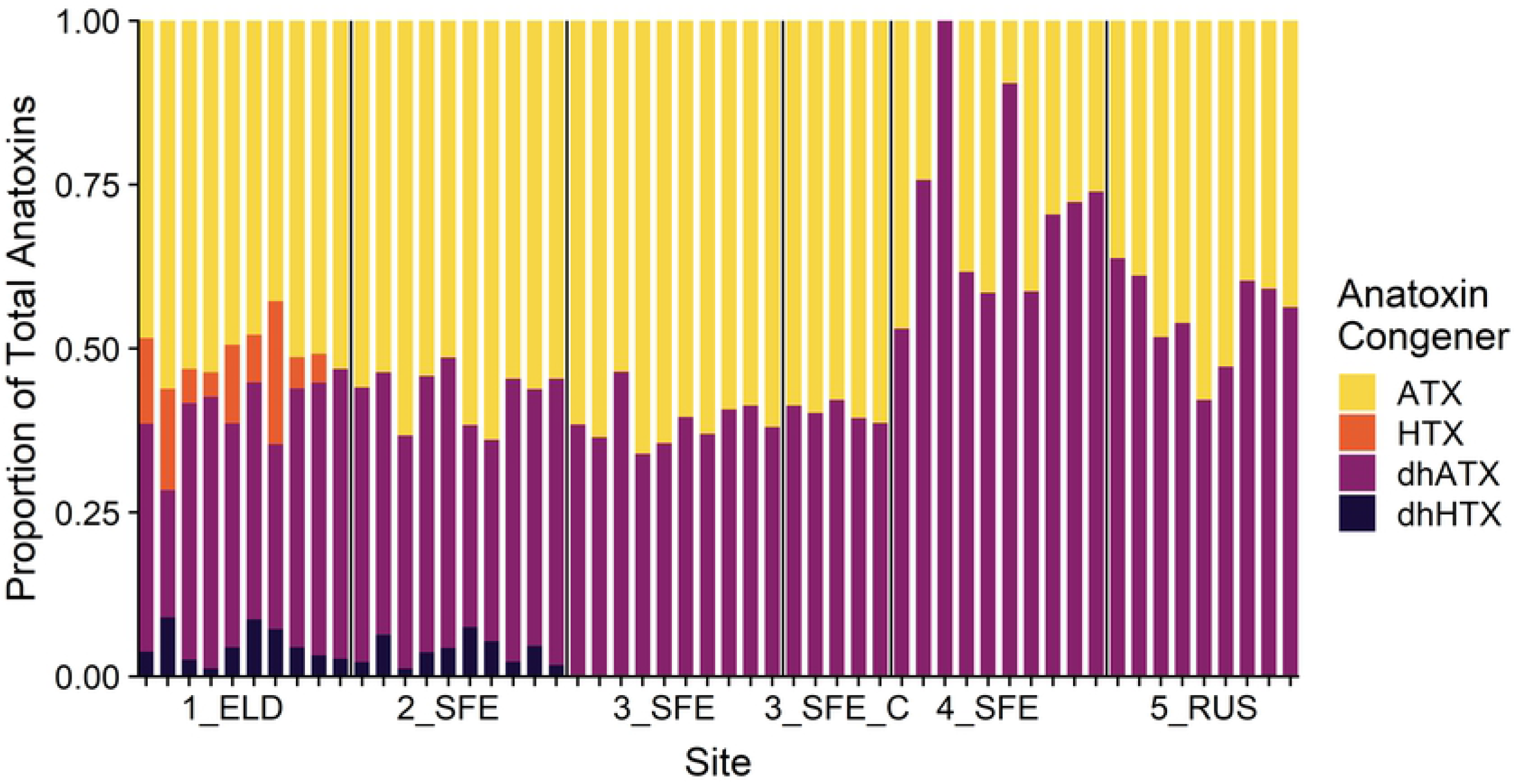
The composition of anatoxin congeners from attached periphyton samples collected at the five sites. Each vertical bar represents one periphyton sample and the vertical black lines delineate each site. 3_SFE_C represents *Cladophora*-dominated samples at site 3_SFE. ATX = anatoxin-a, HTX = homoanatoxin-a, dhATX = dihydroanatoxin-a, dhHTX = dihydrohomoanatoxin-a.

### 3.2 Potential cyanotoxin producers

At sites 3_SFE, 4_SFE and 5_RUS the PCR reactions were positive for *anaC*, but not at sites 1_ELD and 2_SFE. All the *anaC* sequences were identical and most closely matched those of *Oscillatoria* sp. from the Pasteur Culture Collection (*Oscillatoria* sp. PCC 10601, PCC 9240; GenBank accession: JF803652, JF803653; 100% cover and 99.7% identity) and *Phormidium autumnale* (now *Microcoleus autumnalis*; CAWBG618; GenBank accession: KX016036; 93% cover and 99.6% identity) from the Cawthron Institute Culture Collection of Microalgae (http://cultures.cawthron.org.nz/). Comparison of sequences from this study to draft metagenomes from Bouma-Gregson et al. [18] revealed 100% nucleotide sequence identity with the *anaC* gene from the draft *M. autumnalis* genomes assembled in their study. The closest *Anabaena* sp. sequence using BLASTn (*Anabaena circinalis*, GenBank accession: JF803647) yielded only 84.4% nucleotide sequence identity. Samples from site 3_SFE had PCR detections for the *mcyE/nduF* genes. Sequences from these reactions were identical and shared the closest sequence similarity to *Nodularia spumigena* (GenBank accession: CP020114; 100% cover and 99.7% identity). All sequences obtained from the attached and floating samples (*n* = 3) of *C. glomerata* were identical to the above sequences from the cyanobacteria-dominated samples. Saxitoxin and cylindrospermopsin genes were not detected in any of the samples.

### 3.2 Within-site and within-mat cyanotoxin variability

Three sites were assessed for within site variability (3_SFE, 4_SFE and 5_RUS). Low levels of variability in anatoxin concentrations (< 5-fold) were observed within each site (Fig 4A). When anatoxin concentrations were normalised to the concentration of *anaC* gene copies (Fig 4B), the variability was reduced to < 2-fold in each site (Fig 4C). Where samples had high anatoxin concentrations relative to others from the same site (e.g., 5_RUS sample 5; Fig 4), normalisation to the number of toxic cells reduced the anatoxin quota to levels comparable with the other samples. The mean anatoxin quota of 3_SFE was 0.98 pg cell^−1^, while 4_SFE and 5_RUS had quotas of 0.23 pg cell^−1^ and 0.12 pg cell^−1^, respectively. The mean anatoxin quota among all three sites were significantly different (*p* < 0.05; Kruskal-Wallis and Wilcoxon tests).

**Fig 4.**
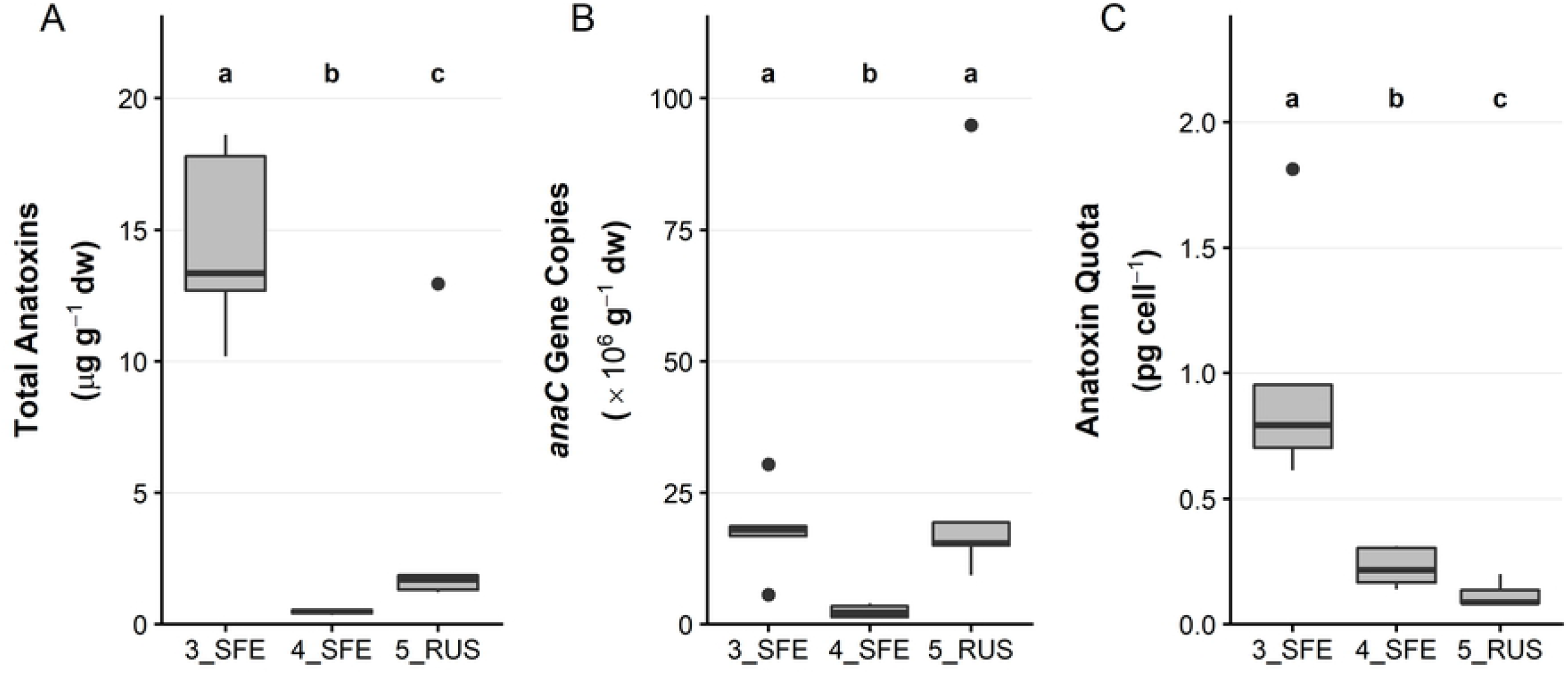
Anatoxin concentrations, *anaC* gene copies and anatoxin quota among three sites on the Eel and Russian rivers. (A) Summed anatoxin concentrations; (B) *anaC* gene copies; and (C) anatoxin quota, from five attached periphyton samples collected from two sites on the Eel River and one site on the Russian River. dw = dried weight. Note the different *y*-axis scales. See Fig 2 for interpretation of boxplots. A Kruskal-Wallis test and pairwise Wilcoxon rank sum test with a Benjamini-Hochberg adjustment was used to identify sites that were significantly different from one another (p <0.05), denoted by the letter above the plot.

Among the five within-mat samples collected from each of three rocks at 4_SFE, total anatoxins (the sum of the four congeners) per mg dw varied by 7-fold. There were no obvious patterns among mats and no significant differences between anatoxin gene copies or quota (*p* > 0.05; Fig 5). Individual samples with high anatoxin concentrations had comparable anatoxin quota upon normalisation to the abundance of toxic cells.

**Fig 5.**
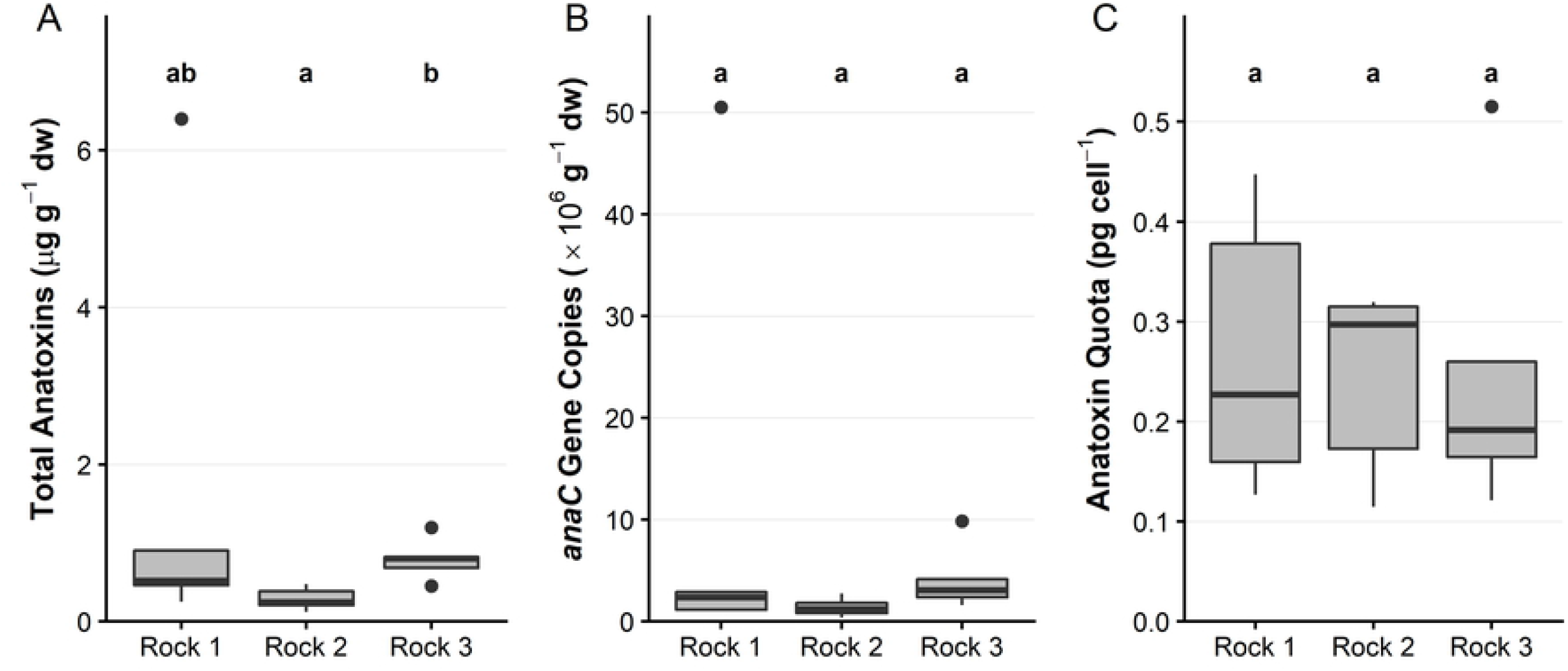
Anatoxin concentrations, anaC gene copies and anatoxin quota among fine-scale samples from three rocks at 4_SFE on the Eel river. (A) Summed anatoxin concentrations; (B) *anaC* gene copy concentrations; and (C) anatoxin quota from five mat samples collected from each of three rocks at 4_SFE. dw = dried weight. Note the different *y*-axis scales. See Fig 2 for interpretation of boxplots. A Kruskal-Wallis test and pairwise Wilcoxon rank sum test with a Benjamini-Hochberg adjustment was used to identify rocks that were significantly different from one another (p <0.05), denoted by the letter above the plot.

## 4. Discussion

### 4.1 Cyanotoxin producers

The first aim of this study was to use molecular techniques to identify potential cyanotoxin producers in benthic mats from the Eel and Russian rivers. The *anaC* gene sequences strongly indicate that *Microcoleus* is the anatoxin producer, both in the *Microcoleus* and *Anabaena*-dominated mat samples. There was no evidence that *Anabaena* was an anatoxin producer, with the *anaC* gene nucleotide sequence identity less than 85% between our sequences and the closest *Anabaena* sp. match. This is consistent with results from *Anabaena* cultures isolated, and *Anabaena* draft-genomes assembled from the Eel and Russian rivers (unpublished observations; K.B.G.). This study detected the *anaC* gene at sites where Bouma-Gregson et al. [18] did not find anatoxin biosynthesis gene clusters in their metagenomes; however our results correspond with previous anatoxin detections in the upper reaches of the Eel watershed [16]. Bouma-Gregson et al. [18] found that only one of four *Microcoleus* strains detected in the metagenomes contained the anatoxin biosynthesis gene cluster. The lack of *anaC* detections with the ddPCR at 1_ELD and 2_SFE despite anatoxins being present at very low concentrations likely reflects the difficulty in completely homogenising samples and highlights the sensitivity of LC-MS/MS as an analytical detection method for these toxins.

Sequencing identified a nodularin producer, with the gene most closely resembling that from *N. spumigena*. This species is generally found in brackish water, though it has been reported in freshwater lakes in Europe based on morphometric identification [27]. Culturing of the causative species and sequencing of other genes involved in nodularin biosynthesis is required to determine whether a novel nodularin producer is present. Sequences for *mcyE* were not detected in the samples, likely due to low abundances of *mcyE* genes compared with *ndaF* genes. Despite this, the presence of microcystins indicates that a yet-to-be-identified microcystin producer is also present in the Eel River.

### 4.2 Concentrations and variability in anatoxin, microcystin and nodularin

The concentrations of anatoxins, NOD-R and MCYs were consistent with data in previous studies from these sites [16, 28], with anatoxin concentrations five times greater than microcystin/nodularin concentrations. Solid-phase adsorption toxin tracking samplers (SPATTs) in the Eel River accumulated ATX and MCYs in 53% – 54% and 41% – 76% of samples for each toxin, respectively [16]. The site with the highest concentrations of both anatoxins and NOD-R was dominated by *Anabaena*, a finding that contrasts with that of Bouma-Gregson et al. [16], who found no differences in ATX concentrations between *Anabaena* and *Microcoleus* dominated mats, although the authors only measured ATX, not dhATX so may have underestimated the total anatoxin concentrations. This difference also demonstrates the important inter-annual variability of periphyton proliferations which highlights the need to further understand within mat, within site, within season and inter-annual variation.

All samples were dominated by ATX and dhATX, and not HTX and dhHTX, which contrasts with observations elsewhere: dhATX, dhHTX and HTX typically dominate in New Zealand [8, 15], and in France ATX dominates [29], although dhATX has also been detected [30]. However, Anderson et al. [19] used anatoxins in crude extracts of *Microcoleus* cultures isolated from the Russian river in their toxicological study and dhATX comprised > 99% of the anatoxins, demonstrating that there is likely large strain to strain variability in the variants produced. The high proportion of dhATX highlights the need to use analytical methods that detect this congener (along with HTX and dhHTX) to prevent underestimation of the risk posed by benthic cyanobacteria in these systems.

Spatial variability of anatoxin content occurred both among and within sites. The magnitude of this variability was relatively low, and upon normalisation to anatoxin quotas, the variability was reduced, a result that is consistent with other studies on *Microcoleus*-dominated mat samples [15, 20]. Anatoxin quotas in this study (0.12 pg cell^−1^ – 0.98 pg cell^−1^) were similar to those observed in the Cardrona River in New Zealand [mean 0.44 pg cell^−1^; 15], however, anatoxin quotas in excess of 7.5 pg cell^−1^ have been reported [20]. Anatoxin quotas can vary both within and between rivers, with 16-to 42-fold differences in anatoxin quotas observed for samples collected at the same site on the same day [15]. Variability in the anatoxin quotas observed in *Microcoleus*-dominated mats may be the result of different toxic genotypes with varying capacities of toxin production [15, 31]. Our results, similarly, indicate that anatoxin concentrations in benthic cyanobacterial mats from the Eel and Russian rivers are largely driven by the abundance of toxic cells. The low variability in anatoxin quotas at both site and within-rock scales observed in this study may, therefore, result from lower diversity in anatoxin-producing strains in the Eel and Russian rivers. Isolation and culture of *Microcoleus* strains would enable the characterisation of toxic genotypes in order to confirm this.

### 4.3 Cyanotoxins and producers in green alga dominant mats

Mats of the green alga *C. glomerata* at 3_SFE yielded concentrations of anatoxins and NOD-R comparable with those found in the cyanobacteria-dominated samples. *Cladophora glomerata* is globally distributed and supports complex epiphytic algal and microbial assemblages [11]. Previous analysis of similar mats from these rivers has shown the presence of a variety of cyanobacterial taxa including Oscillatoriacae [10]. The presence of cyanotoxins at such high concentrations in the *C. glomerata*-dominated mats in this system raises important questions about how extensive this phenomenon is in other non-cyanobacterial dominated mats. Further studies of cyanotoxins in periphyton mats should be a priority for future research to assist with increasing knowledge on the risks associated with cyanotoxins in river systems.

### 4.3 Conclusions

The data from this study provides further evidence that *Microcoleus* and *Nodularia* are anatoxin- and nodularin-producers in the Eel and Russian rivers and enables targeted culturing of these taxa in future studies for definitive confirmation. Our data also show that when these organisms are not macroscopically visible, they can still produce toxin concentrations equivalent to those mats where they are macroscopically dominant. The spatial variability in toxin concentrations among samples highlights the need for careful sampling design when assessing toxin levels at a site for risk assessment purposes. Normalisation of anatoxin concentration data to quotas reduced variability among samples indicating that a key driver in toxin variability is the abundance of toxic genotypes present in a sample. Cyanotoxins were present at concerning concentrations in mats dominated by the green alga *C. glomerata*. Further research to investigate the extent of this phenomenon should be prioritised.

## Acknowledgements

This research was performed at the UCNRS Angelo Coast Range Reserve. We would like to especially thank the reserve manager, Peter Steel, for his assistance. We also thank Mary Powers (UC, Berkeley) for her assistance in the field.

## Supplementary Information

**S1 Table.** Physicochemical parameters collected at each sampling site. DO = dissolved oxygen. DIN = dissolved inorganic nitrogen. DRP = dissolved reactive phosphorus.

**S2 Table.** Primers and probes used for PCR screening for cyanotoxin genes and for ddPCR analysis of *anaC* gene copy number.

**S1 Fig.** Samples collected from the five sites were analysed for toxins, anatoxin quota and within-rock variation in anatoxin-quota. Squares in the grids represent 1 m^2^. Fine-scale samples consisted of five samples collected from periphyton on a single cobble.

**S3 Table.** The mean proportion of each anatoxin congener from attached periphyton samples collected at the five sites ± standard deviation (n = 10 for all sites except 5_RUS where n = 9).

## Literature Cited

1. Edwards C, Beattie KA, Scrimgeour CM, Codd GA. Identification of anatoxin-a in benthic cyanobacteria (blue-green algae) and in associated dog poisonings at Loch Insh, Scotland. Toxicon. 1992;30(10):1165–75.

2. Mez K, Beattie KA, Codd GA, Hanselmann K, Hauser B, Naegeli H, et al. Identification of a microcystin in benthic cyanobacteria linked to cattle deaths on alpine pastures in Switzerland. Eur J Phycol. 1997;32(2):111–7.

3. Hamill KD. Toxicity in benthic freshwater cyanobacteria (blue-green algae): First observations in New Zealand. New Zealand Journal of Marine and Freshwater Research. 2001;35(5).

4. Hudon C, De Sève M, Cattaneo A. Increasing occurrence of the benthic filamentous cyanobacterium *Lyngbya wollei*: a symptom of freshwater ecosystem degradation. Freshwater Science. 2014;33(2):606–18.

5. Gaget V, Humpage AR, Huang Q, Monis P, Brookes JD. Benthic cyanobacteria: A source of cylindrospermopsin and microcystin in Australian drinking water reservoirs. Water Research. 2017;124:454–64.

6. Belykh OI, Tikhonova IV, Kuzmin AV, Sorokovikova EG, Fedorova GA, Khanaev IV, et al. First detection of benthic cyanobacteria in Lake Baikal producing paralytic shellfish toxins. Toxicon. 2016;121:36–40.

7. Quiblier C, Wood S, Echenique-Subiabre I, Heath M, Villeneuve A, Humbert J-F. A review of current knowledge on toxic benthic freshwater cyanobacteria–ecology, toxin production and risk management. Water Research. 2013;47(15):5464–79.

8. McAllister TG, Wood SA, Hawes I. The rise of toxic benthic *Phormidium* proliferations: A review of their taxonomy, distribution, toxin content and factors regulating prevalence and increased severity. Harmful Algae. 2016;55:282–94. doi: 10.1016/j.hal.2016.04.002.

9. Cantoral Uriza E, Asencio A, Aboal M. Are we underestimating benthic cyanotoxins? Extensive sampling results from Spain. Toxins. 2017;9(12):385. doi:10.3390/toxins9120385.

10. Furey PC, Lowe RL, Power ME, Campbell-Craven AM. Midges, *Cladophora*, and epiphytes: shifting interactions through succession. Freshwater Science. 2012;31(1):93–107.

11. Zulkifly S, Hanshew A, Young EB, Lee P, Graham ME, Graham ME, et al. The epiphytic microbiota of the globally widespread macroalga *Cladophora glomerata* (Chlorophyta, Cladophorales). American Journal of Botany. 2012;99(9):1541–52.

12. Merel S, Villarín MC, Chung K, Snyder S. Spatial and thematic distribution of research on cyanotoxins. Toxicon. 2013;76:118–31.

13. Kaebernick M, Neilan BA. Ecological and molecular investigations of cyanotoxin production. FEMS Microbiol Ecol. 2001;35(1):1–9.

14. Pearson LA, Neilan BA. The molecular genetics of cyanobacterial toxicity as a basis for monitoring water quality and public health risk. Current Opinion in Biotechnology. 2008;19(3):281–8. doi: 10.1016/j.copbio.2008.03.002.

15. Wood S, Puddick J. The abundance of toxic genotypes is a key contributor to anatoxin variability in *Phormidium*-dominated benthic mats. Marine Drugs. 2017;15(10):307. doi:10.3390/md15100307.

16. Bouma-Gregson K, Kudela RM, Power ME. Widespread anatoxin-a detection in benthic cyanobacterial mats throughout a river network. PLOS ONE. 2018;13(5):e0197669. doi: 10.1371/journal.pone.0197669.

17. Fetscher AE, Howard MD, Stancheva R, Kudela RM, Stein ED, Sutula MA, et al. Wadeable streams as widespread sources of benthic cyanotoxins in California, USA. Harmful Algae. 2015;49:105–16.

18. Bouma-Gregson K, Olm MR, Probst AJ, Anantharaman K, Power ME, Banfield JF. Impacts of microbial assemblage and environmental conditions on the distribution of anatoxin-a producing cyanobacteria within a river network. The ISME Journal. 2019. doi: 10.1038/s41396-019-0374-3.

19. Anderson B, Voorhees J, Phillips B, Fadness R, Stancheva R, Nichols J, et al. Extracts from benthic anatoxin-producing *Phormidium* are toxic to 3 macroinvertebrate taxa at environmentally relevant concentrations. Environmental Toxicology and Chemistry. 2018;37(11):2851–9.

20. Kelly L, Wood S, McAllister T, Ryan K. Development and application of a quantitative PCR assay to assess genotype dynamics and anatoxin content in *Microcoleus autumnalis*-dominated mats. Toxins. 2018;10(11):431. doi:10.3390/toxins10110431.

21. Wood SA, Puddick J, Fleming R, Heussner AH. Detection of anatoxin-producing *Phormidium* in a New Zealand farm pond and an associated dog death. New Zealand Journal of Botany. 2017;55(1):36–46.

22. Wood SA, Kuhajek JM, de Winton M, Phillips NR. Species composition and cyanotoxin production in periphyton mats from three lakes of varying trophic status. FEMS Microbiol Ecol. 2012;79(2):312–26.

23. Jungblut A-D, Neilan BA. Molecular identification and evolution of the cyclic peptide hepatotoxins, microcystin and nodularin, synthetase genes in three orders of cyanobacteria. Arch Microbiol. 2006;185(2):107–14.

24. Ballot A, Fastner J, Wiedner C. Paralytic shellfish poisoning toxin-producing cyanobacterium *Aphanizomenon gracile* in Northeast Germany. Applied and Environmental Microbiology. 2010;76(4):1173–80. doi: 10.1128/aem.02285-09.

25. Rantala-Ylinen A, Känä S, Wang H, Rouhiainen L, Wahlsten M, Rizzi E, et al. Anatoxin-a synthetase gene cluster of the cyanobacterium *Anabaena* sp. strain 37 and molecular methods to detect potential producers. Applied and Environmental Microbiology. 2011;77(20):7271–8.

26. Mihali TK, Kellmann R, Muenchhoff J, Barrow KD, Neilan BA. Characterization of the gene cluster responsible for cylindrospermopsin biosynthesis. Applied and Environmental Microbiology. 2008;74(3):716–22. doi: 10.1128/aem.01988-07.

27. Akcaalan R, Mazur-Marzec H, Zalewska A, Albay M. Phenotypic and toxicological characterization of toxic *Nodularia spumigena* from a freshwater lake in Turkey. Harmful Algae. 2009;8(2):273–8.

28. Bouma-Gregson K, Power ME, Bormans M. Rise and fall of toxic benthic freshwater cyanobacteria (*Anabaena* spp.) in the Eel river: Buoyancy and dispersal. Harmful Algae. 2017;66:79–87.

29. Echenique-Subiabre I, Tenon M, Humbert J-F, Quiblier C. Spatial and temporal variability in the development and potential toxicity of *Phormidium* biofilms in the Tarn River, France. Toxins. 2018;10(10):418. doi:10.3390/toxins10100418.

30. Mann S, Cohen M, Chapuis-Hugon F, Pichon V, Mazmouz R, Méjean A, et al. Synthesis, configuration assignment, and simultaneous quantification by liquid chromatography coupled to tandem mass spectrometry, of dihydroanatoxin-a and dihydrohomoanatoxin-a together with the parent toxins, in axenic cyanobacterial strains and in environmental samples. Toxicon. 2012;60(8):1404–14.

31. Wood SA, Smith FM, Heath MW, Palfroy T, Gaw S, Young RG, et al. Within-mat variability in anatoxin-a and homoanatoxin-a production among benthic *Phormidium* (cyanobacteria) strains. Toxins. 2012;4(10):900–12.

